# An age-specific burial practice reflected in ancient DNA preservation in Neolithic Çatalhöyük

**DOI:** 10.1101/2024.12.13.628343

**Authors:** Ayça Küçükakdağ Doğu, Fatma Küçük Baloğlu, Maciej Chyleński, Eline M.J. Schotsmans, Muhammed Sıddık Kılıç, Kıvılcım Başak Vural, Eren Yüncü, Damla Kaptan, Merve N. Güler, Sevgi Yorulmaz, Nergis Bilge Karabulut, Ekin Sağlıcan, Duygu Deniz Kazancı, Marco Milella, Arkadiusz Marciniak, Yılmaz Selim Erdal, Marin Pilloud, Clark Spencer Larsen, Anders Götherström, Ian Hodder, Füsun Özer, Scott D. Haddow, Christopher J. Knüsel, Mehmet Somel

## Abstract

Selective funerary practices can inform about social relationships in prehistoric societies but are often difficult to discern. Here we present evidence for an age-specific practice at the Neolithic site of Çatalhöyük in Anatolia, dating to the 7th millennium BCE. Among ancient DNA libraries produced from 362 petrous bone samples, those of subadults contained three times higher average human DNA than those of adults. This difference in organic preservation was also confirmed by FTIR analysis. Studying similar datasets from seven prehistoric and historical sites, we found a similar age-related difference in only one cemetery. We propose that the organic preservation difference with age was caused by the special treatment of chosen corpses before interment, such as defleshing or drying, which was more frequently applied to Çatalhöyük adults and promoted organic decay.

## Results and Discussion

The early Holocene Neolithic transition in Southwest Asia to sedentism and agriculture was accompanied by the adoption of elaborate and standardized funerary practices, including widespread intramural burials, the curation of and defleshing of bodies before burial, as well as the removal and plastering of skulls and their reburial in skull caches (Boz and Hager 2008; Kuijt 2002; Rollefson 1986; Özdoğan and Özdoğan 1989; Benz 2009; Belfer-Cohen and Goring-Morris 2011; Erdal 2015; Bocquentin, Kodas, and Ortiz 2016).

Decades of excavations at the Central Anatolian Neolithic settlement of Çatalhöyük have revealed an abundance of such funerary practices. Çatalhöyük East Mound was a relatively large Pottery Neolithic site occupied continuously between 7100-5950 BCE (Bayliss et al. 2015), with estimates ranging between 500 and 5000 inhabitants (Cessford 2001; Kuijt and Marciniak 2024). The settlement is known for its dense housing, wall paintings, striking imagery and figurines (Ian Hodder and Cessford 2004; Ian Hodder 2021; I Hodder 2007). About 700 human skeletal remains, either as complete individuals in their original location (primary burials), or as burials moved to another location (secondary) or loose skeletal elements (tertiary) have been excavated to date; these are mostly individual burials, with roughly even numbers of subadults and adults, and also similar numbers of females and males (Larsen et al. 2019; Knüsel et al. 2021; Haddow et al. 2021). These burials revealed a number of funerary patterns. One involves burial location, with a higher frequency of adult burials under the well-kept platform areas compared to a higher frequency of subadult burials near the oven or in side-rooms (Boz and Hager 2008). Large numbers of bodies showed evidence of binding and extreme flexion (Pilloud et al. 2016; Knüsel et al. 2021); many were deposited with pigments (Schotsmans et al. 2022) and burial objects, such as pottery and stone tools (Vasic, Knüsel, and Haddow 2021). About 3% of the burials were found with their complete skull (cranium and mandible) removed, along with 5% of crania found separated; the curation of collected skulls with plastering and pigment is also well-documented (Haddow and Knüsel 2017). Another pattern that emerges is the increase in secondary and tertiary burials over the site’s occupation (Haddow et al. 2021).

We had previously reported a trend towards higher aDNA preservation in juvenile temporal bones versus adult temporal bones at Çatalhöyük (Yaka et al. 2021), raising the question of whether aDNA preservation levels may be related to age-specific burial treatments. We recently generated a larger aDNA dataset comprising the skeletal remains of 395 individuals as part of an investigation into the social structure of Çatalhöyük, the results of which may be found in an accompanying study (Yüncü et al. 2024). Here, we present findings from this dataset on age-related differences in organic preservation in Çatalhöyük human remains and in other archaeological sites with published aDNA datasets.

We combined all shotgun-sequenced aDNA libraries obtained from 362 Çatalhöyük individuals, restricting our analysis only to libraries produced from petrous bones (Supplementary Methods). The median endogenous human aDNA proportion across these libraries was only 0.45%. This is a low value but is expected given the age of the site and the relatively temperate climatic conditions of the region. However, the median endogenous human DNA proportions of subadults (n=230, prenatal to adolescent individuals; Supplementary Methods) was 0.68%, nearly three times higher than the adult proportion of 0.24% adults (n=132) [Mann-Whitney U (MWU) test, two-sided p=3e-7]. Separating the sample into further age subcategories revealed a clear monotonic decline in endogenous proportion with individual age, with an approximately four times difference in median endogenous human DNA proportions between prenatal individuals and adults (Spearman correlation r=-0.300, two-sided p=6e-9) **(Figure 1A)**. Notably, adolescents had values similar to those of adults, while children appeared intermediate.

**Figure 1:**
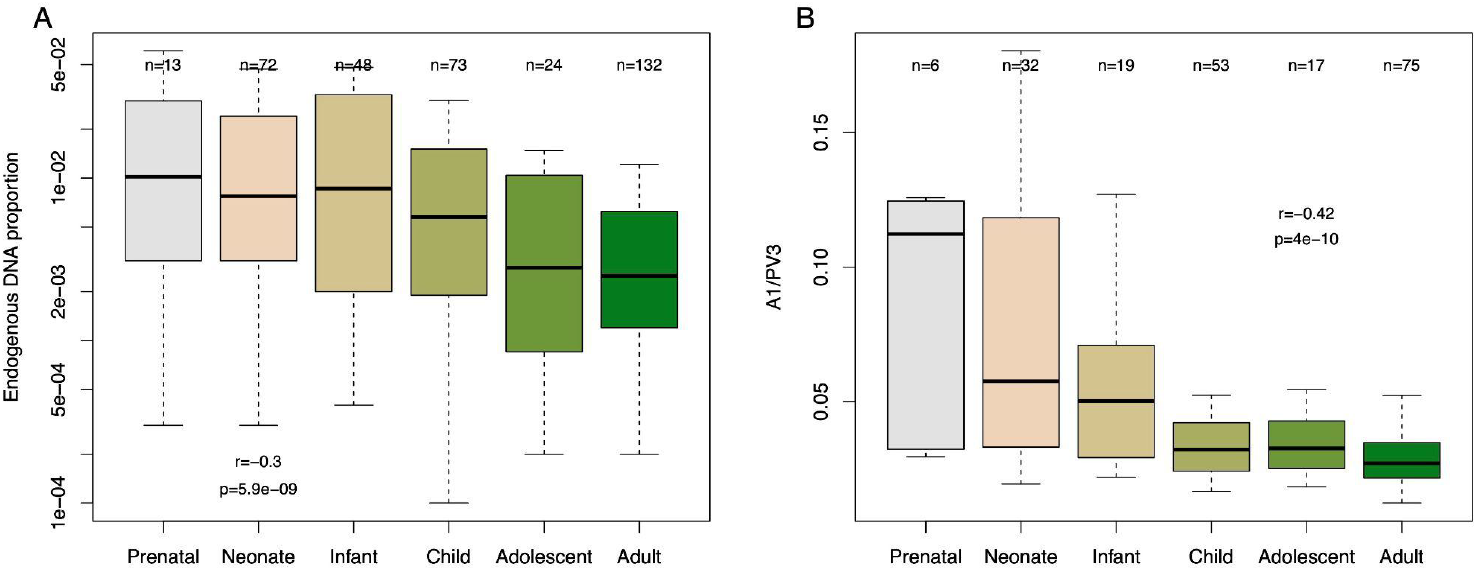
DNA preservation and organic preservation measured using FTIR spectra DNA in Çatalhöyük burials. **A)** Endogenous human DNA yield across libraries of 362 individuals. All libraries were produced using petrous bone samples. We used the raw human DNA proportion (before removing duplicates) and only libraries from petrous bones. **B)** Amide I (A1)/Phosphate (PV3) ratios of FTIR spectra measured from petrous bone samples in 202 individuals. The values plotted on the top show sample sizes. The r and p values are calculated using the Spearman correlation test. For a definition of age categories, see Supplementary Methods. The boxplots are plotted without outliers for visual clarity.

There was no reason to suspect batch effects for this change because skeletal samples were not processed based on age. Still, to exclude that the age effect might be caused by technical issues in DNA extraction or library preparation, we investigated organic preservation using an independent method, Attenuated Total Reflectance - Fourier Transform Infrared Spectroscopy (ATR-FTIR), which can measure organic preservation in archaeological bone (Leskovar et al. 2020; Kontopoulos et al. 2020). We compared the ratio of Amide 1 (A1) and Phosphate 3 (PV3) FTIR-ATR spectra across 202 Çatalhöyük bone samples (Supplementary Methods). This revealed the same age-related decline pattern, suggesting the effect involves organic preservation of the bone and is not particular to aDNA (Spearman correlation r=-0.42, two-sided p=4e-10) (**Figure 1B; Table_S1**).

We asked if the reason for the observed age difference in organic preservation at Çatalhöyük could be linked to age-related changes in the density of the petrous bone, e.g. by remodeling (Kontopoulos et al. 2019), which could influence organic preservation. If so, the same pattern should be detectable in other paleogenome datasets. We studied endogenous DNA in published aDNA libraries dating to between 10,000 to 1000 years ago with sufficient sample sizes, restricting the analysis to those produced from petrous bones. Our dataset included intramural burials in an Upper Mesopotamian Pre-Pottery Neolithic village (Altinişik et al. 2022) from c.8500-7500 BCE, (Çayönü, n=29), burials in three Neolithic Central European sites (Gelabert et al. 2024) and one West European site (Rivollat et al. 2023) dating to c.5500-4500 BCE, including within-settlement burials or burials in cemeteries (Polgár-Ferenci-hát, n=43; Asparn-Schletz, n=55; Nitra Horné Krškany, n=36; Gurgy, n=94), and burials in three Avar cemeteries from Early Medieval Central Europe dating to c.500-800 CE (Gnecchi-Ruscone et al. 2024) (Kunszállás, n=44, Rákóczifalva, n=214). Only one of the seven cemeteries tested, Rakoczifalva used by the early Medieval Avars, showed a significant trend towards higher endogenous human DNA in subadults as at Çatalhöyük (MWU test, two-sided p=0.008) (**Figure 2**). Conversely, burials in one Central European Neolithic cemetery showed a significantly higher endogenous human DNA in adults (MWU test, two-sided p=0.01). Data from the other five sites showed no significant trends and a range of tendencies.

**Figure 2:**
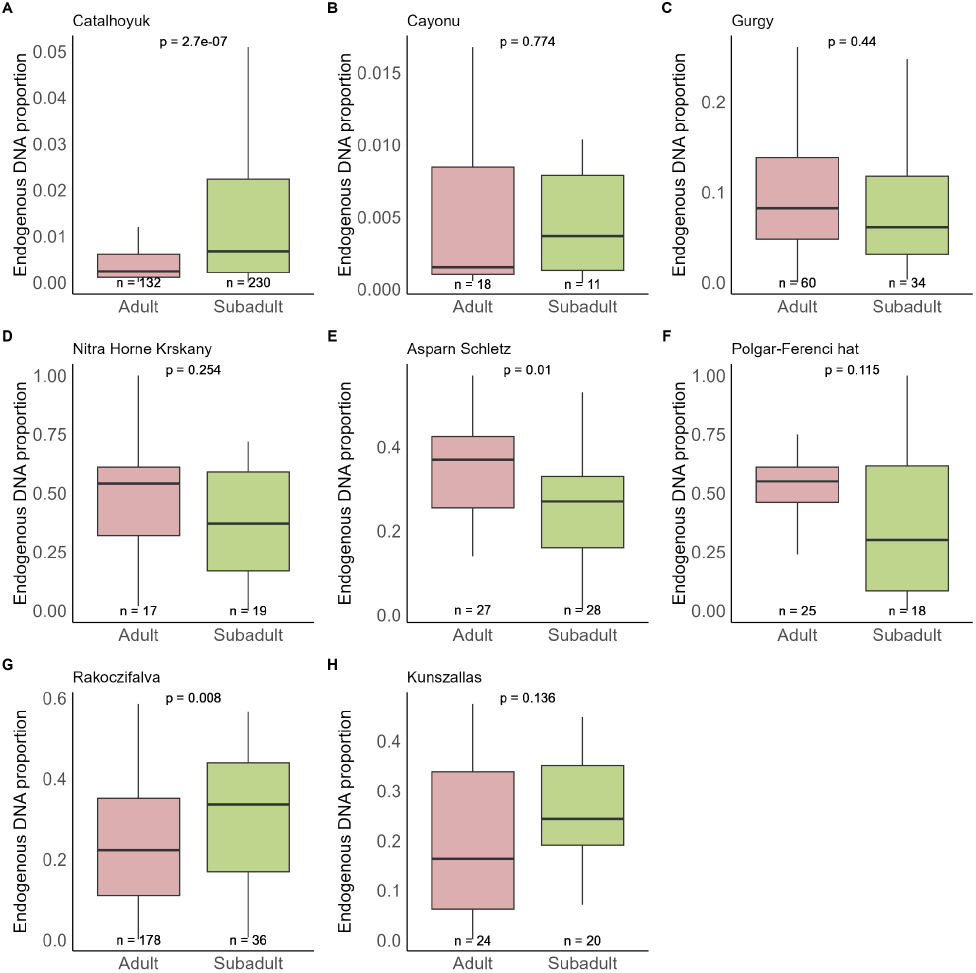
DNA preservation in Çatalhöyük **(A)** and in published datasets from early Holocene Anatolian Çayönü (**B**), Gurgy from mid-Holocene West Europe, (**C**), Polgár-Ferenci-hát, Asparn-Schletz, and Nitra Horné Krškany from mid-Holocene Central Europe (**D-F**), and Kunszállás and Rákóczifalva from early Medieval Central Europe (**G-H**). We only used libraries that were produced using petrous bone samples (or “cochlea”). Individuals with no age information or ambiguous categorization (e.g. “juvenile/adult”) were not included. We used the raw human DNA proportions reported by the respective authors. The values plotted on the bottom show sample sizes. The p values are calculated using the MWU test (two-sided). The boxplots are plotted without outliers and each has different y-axis scales for visual clarity.

These results suggest that the age effect on DNA preservation observed at Çatalhöyük is not a universal phenomenon. Instead, the Çatalhöyük pattern could reflect the outcome of a specific treatment of the corpse or burial practices affecting skeletal preservation, differentially applied to the dead of different ages. We note that the effect was observed across all Çatalhöyük periods **(Figure S1)**, suggesting that this putative practice persisted across centuries.

We sought possible correlates of this differential preservation pattern, asking whether the burials with low versus high preservation of aDNA a) were identified as primary, primary disturbed (indicating the grave being reopened), secondary, or tertiary burials (**Figure S2**), b) showed signs of extreme flexion (**Figure S3**), and c) differed in their burial location (**Figure S4**). Flexion levels had no impact [Kruskal-Wallis (KW) test p=0.12]. Burial type had a modest effect (KW test p=0.013), with a trend for higher endogenous DNA among primary disturbed burials, although the effect was not significant after dividing the data into age groups (KW test p>0.05). Finally, we found a weak association between aDNA preservation and burial location, with neonates buried in platform areas exhibiting lower preservation compared with those from other areas of buildings (KW test p=0.001) (**Figure S1D)**. Still, the effect of burial location was not visible in all age groups, suggesting that the burial location is not the sole reason for the observed difference.

One possible cause of such difference is a pre-burial treatment of selected corpses. This may have included above-ground decomposition, defleshing, drying, and similar processes (Pilloud et al. 2016). Such treatments could have expedited decomposition by boosting microbial activity or compromised bone integrity, leading to lower organic preservation. If this hypothesis is correct, it follows that these procedures were carried out differently based on the age-at-death of individuals: subadults frequently interred directly and adults frequently subject to treatment. An intriguing possibility is that the observed lower DNA preservation of neonates buried under platforms (like adults) (**Figure S4**) could be the result of these specific neonates having undergone an adult-like pre-burial treatment.

An age-biased funerary treatment would also be consistent with similar age differences in skull/cranial removal practices at Çatalhöyük. Haddow and Knüsel report that among 15 skeletons with skulls/crania removed, twelve were adults while only two were adolescents and one was a child, a statistically significant difference (binomial test p=0.014) (Haddow and Knüsel 2017). A similar trend (although weaker) is also seen among cases of isolated crania, where only 5 out of 16 cases belonged to subadults (p=0.067).

Our study contributes to the compendium of age-related practices recorded at Çatalhöyük. It further suggests that aDNA preservation may be used as an indirect indicator of body treatment before interment. Indeed, the pattern of age-related aDNA preservation difference we also find in Early Medieval Avars in Hungary (Gnecchi-Ruscone et al. 2024) raises the question of whether this community may similarly have processed adult bodies in particular ways.

## Acknowledgments

We thank all members of the CompEvo (METU) group and the NEOMATRIX Collective for helpful discussion and suggestions. This work was supported by the H2020 ERC Consolidator grant (no. 772390 NEOGENE to MS), the H2020-WIDESPREAD-05-2020 TWINNING grant (no. 952317 NEOMATRIX to MS), TUBITAK of Turkey (no. 117Z229 to MS and no. 120C217 to FKB), NSF (no. BCS-1827338 to MP). CJK, SDH. EMJS’s participation in this research benefited from the scientific framework of the University of Bordeaux’s IdEx (Initiative d’Excellence) ‘Investments for the Future’ program / GPR (Grands Programmes de Recherche) ‘Human Past’.

## Supplementary Methods

### a. Çatalhöyük genomes

From Çatalhöyük, we combined all libraries from the Neolithic period produced from petrous bone samples (which constitute c.90% of the total). The laboratory and data preprocessing methods are described in (Yaka et al. 2021; Yüncü et al. 2024). Briefly, we pulverized bone material using drilling or cutting, isolated DNA using the Dabney method (Dabney et al. 2013) and produced Meyer-Kircher double-stranded libraries (Meyer and Kircher 2010), which were shotgun sequenced on Illumina HiSeq and NovaSeq platforms at an average of 13 million reads per library. The reads were mapped to hg19 using the *bwa aln* algorithm (Li and Durbin 2009). See the original publications (Yaka et al. 2021; Yüncü et al. 2024) for details.

For libraries was sequenced multiple times, we used the median endogenous percentage value. When calculating the endogenous proportion in ancient DNA libraries, we did not remove duplicate (clonal) molecules because clonality will be affected by the amount of sequencing and PCR cycles. However, we note that many libraries with low preservation have high clonality, and the age differences are larger when using only unique molecules (data not shown).

In downstream analyses, we used osteological age-at-death and the burial context information as defined by the Çatalhöyük Human Remains Team (2009-2017) (Haddow et al. 2021; Knüsel et al. 2021). Age groups were defined as follows: prenatal (<38 weeks’ gestation); neonate (0-2 months); infant (2 months-3 years); child (3-12 years), adolescent (12-20 years), adults (>20 years old).

### b. Published genomic datasets

We compiled age-at-death, tissue type, and endogenous DNA proportion from the four publications cited, available in each publication’s supplementary information files. We limited analyses to petrous bones and removed cases where tissue type or age information was ambiguous.

### c. Fourier Transform Infrared (FTIR) Spectroscopy

We used FTIR analysis for studying organic preservation in Çatalhöyük human skeletal remains, following recent work (Kontopoulos et al. 2020). We selected 202 samples covering six different age categories and sexes [prenatal (n=6), neonate (n=33), infant (n=19), child (n=54), adolescent (n=17) and adult (n=77)]. Note that in Figure 1B we report results for the same 202 individuals included in the genetic analysis in Figure 1A. Sampling was done from the same bone regions used for aDNA analysis. The samples were powdered with the help of a pestle and mortar and convenient sieves were used to standardise the particle size of the powder to 50-100 µm. IR spectra of the samples were collected using a Universal ATR accessory (Specac Ltd., UK) equipped with a Perkin Elmer Spectrum 100 FTIR spectrometer (Perkin-Elmer Inc, Boston, MA, USA). In the ATR-FTIR spectroscopy technique, the atmospheric CO_2_ and H_2_O absorption bands of the environmental air, and a background spectrum were recorded before the sample spectra were collected. This spectrum was mathematically subtracted from the sample spectra. This process was automatically repeated by the software for each sample spectra. Following the subtraction process, sieved sample powder was placed directly onto the diamond crystal platform with slight pressure to ensure that the samples touched the crystal evenly. The scanning process was performed without drying in the 4000 to 450 cm^-1^ spectral range at room temperature, with 64 scans collected at a resolution of 4 cm^-1^ for each spectrum. The spectrum was collected three times independently for each sample. The average spectra of these three replicates were used for further analysis and data manipulation was performed using *OPUS 5.5* software (Bruker Optics, Reinstetten, Germany). All quantitative band intensity calculations were performed using baseline-corrected spectra over the entire spectral range for all samples. We used peak intensity (I) calculations to determine the Amide/Phosphate ratio () which has a strong correlation with organic/mineral content (Trueman et al. 2004; Kontopoulos et al. 2020). Therefore, the I 1638 /I 1014 ratio was used to assess the relative organic content in bones.

## Supplementary Figures

**Figure S1:**
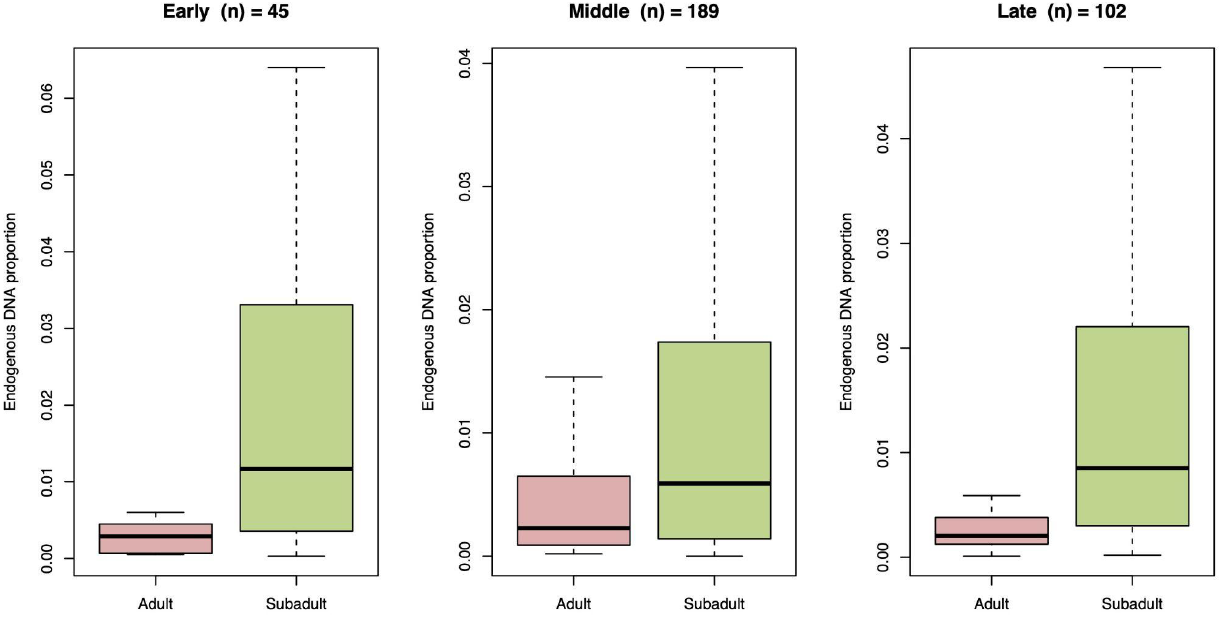
Endogenous human DNA relative to age-at-death in the three main archaeological periods of Çatalhöyük, Early (7100-6700 BCE), Middle (6700-6500 BCE), Late (6500-6300 BCE), and Final (6300-5950 BCE). The age differences in endogenous DNA percentage were significant (MWU test p<0.01) in all cases. The boxplots are plotted without outliers and each has different y-axis scales for visual clarity.

**Figure S2:**
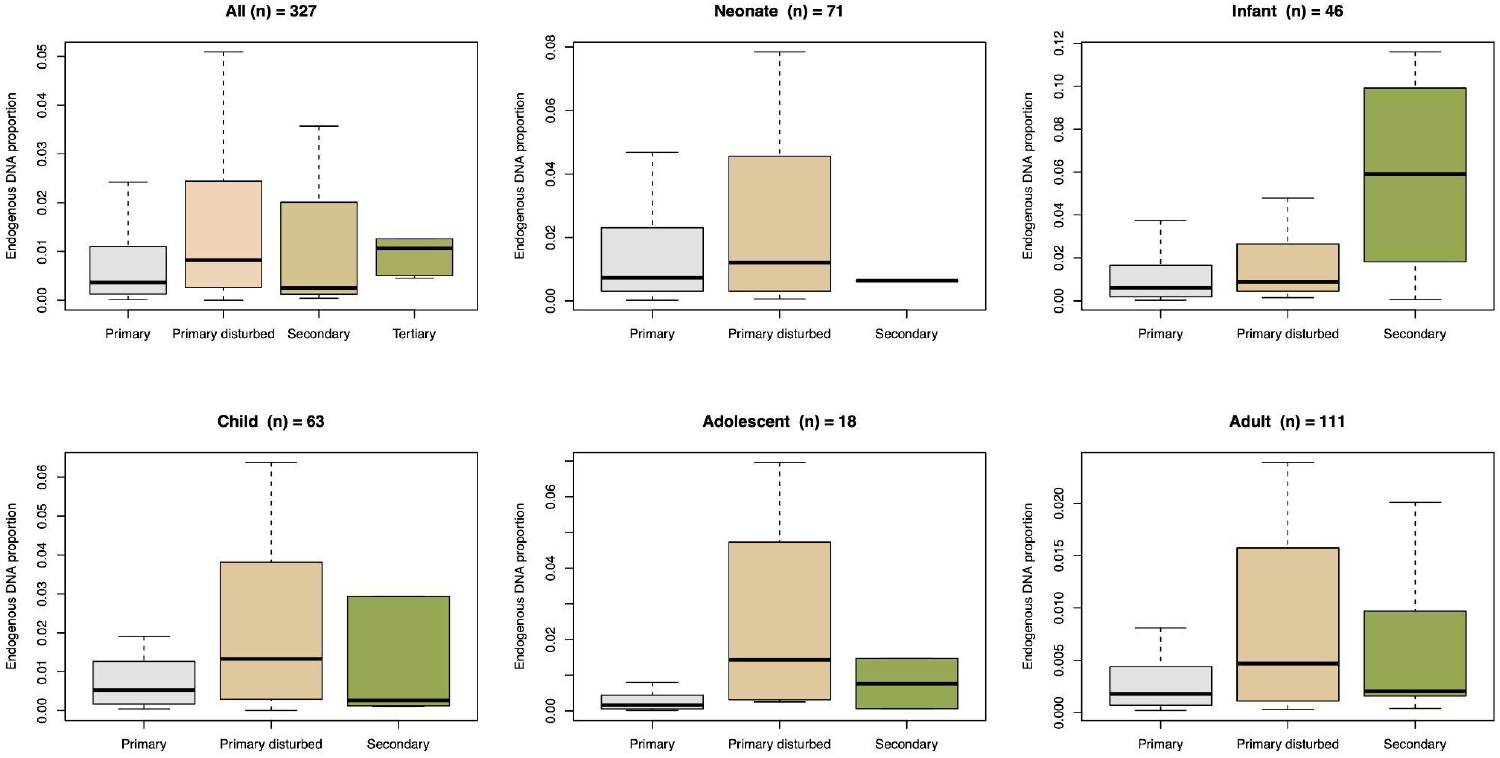
Endogenous human DNA across different burial deposition types, where “Primary” means an undisturbed state, “Primary disturbed” indicates the grave was re-opened, “Secondary” indicates the skeletal remains were removed from the original deposition location, and “Tertiary” indicates remains found outside of burial context. The top left panel shows all the individuals analysed, while the remaining panels show the data divided into age groups. The position type appeared to impact endogenous DNA percentage using all the data (KW test p=0.013). However, the results are not significantly different for separate age groups (KW test p>0.05). The boxplots are plotted without outliers and each has different y-axis scales for visual clarity.

**Figure S3:**
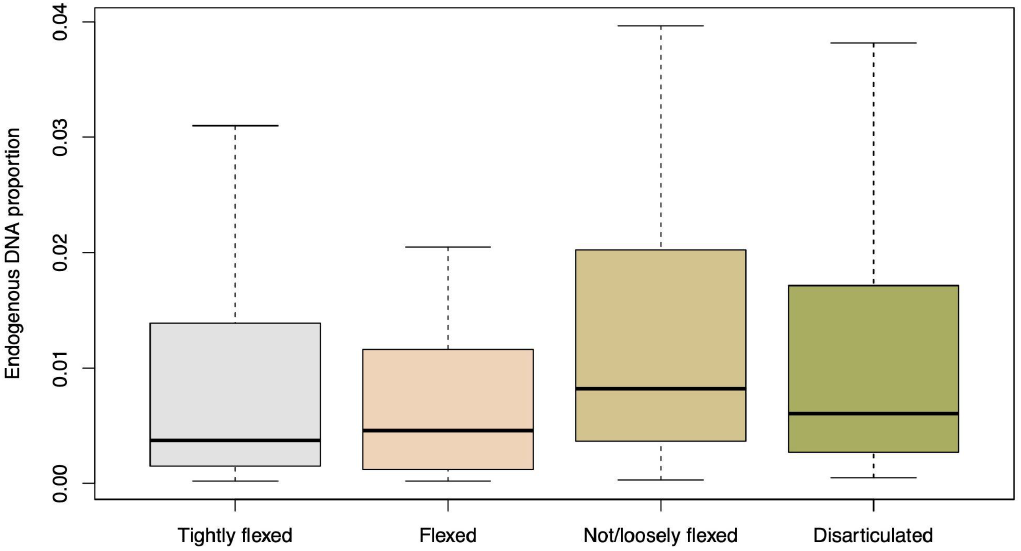
Endogenous human DNA across flexion levels among Çatalhöyük burials. Flexion levels did not show a significant association with endogenous DNA percentage (KW test p=0.12). The boxplots are plotted without outliers for visual clarity.

**Figure S4:**
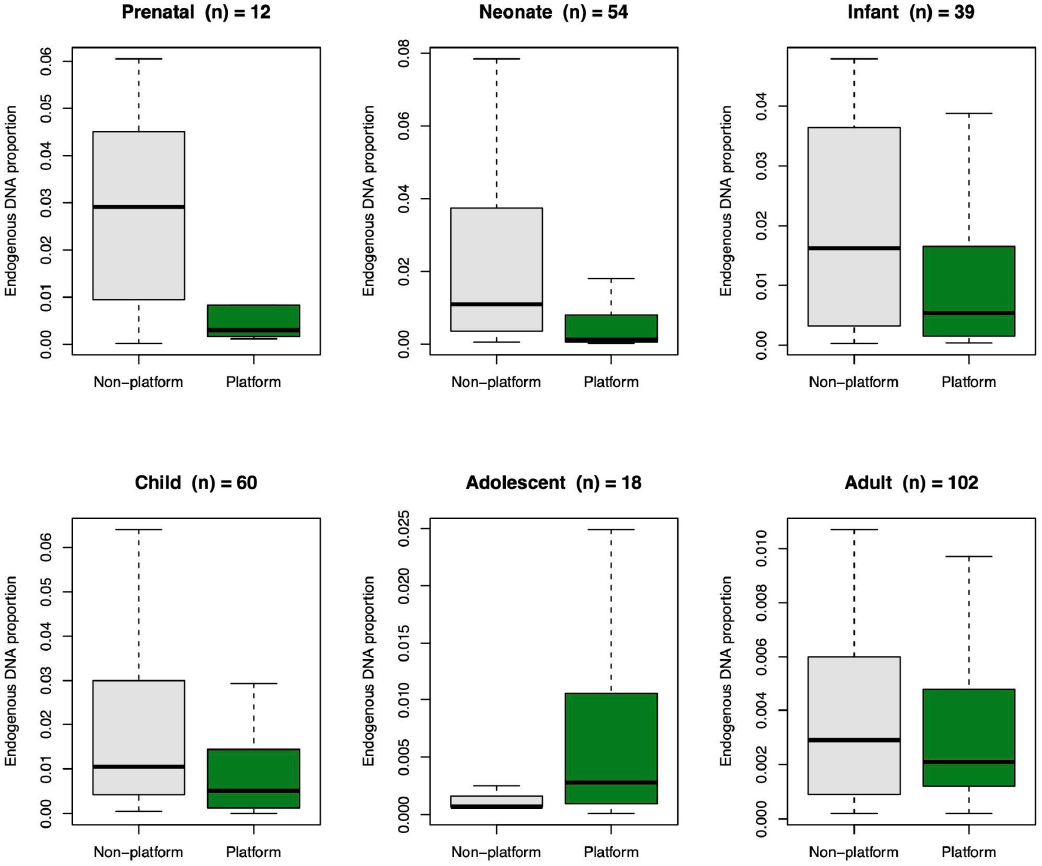
Burial location of all Çatalhöyük individuals within houses, separated by the platform area (mostly limited to adults) and non-platform areas (walls, central floor, corners, or siderooms). The non-platform versus platform distributions were overall significant (MWU test p<0.001). The differences were also significant when limiting the sample to neonates (p=0.006) but not other age groups. The boxplots are plotted without outliers and each has different y-axis scales for visual clarity.

## Notes

### Competing Interest Statement

The authors have declared no competing interest.

